# Induction of hypoxic response despite normoxic conditions is associated with vascular mitochondrial dysfunction in diet induced metabolic syndrome

**DOI:** 10.1101/2025.05.05.652328

**Authors:** Evanka Madan, Vaibhav Jain, Vijay Pal Singh

**Author notes:** Corresponding author; Dr. Evanka Madan, PhD, Consultant and Assistant Professor, Sir Ganga Ram Hospital, Rajinder Nagar, New Delhi, India -110060.

## Abstract

Reduced mitochondrial function is one of the various cellular pathologies in cardio-metabolic syndrome. Elevated asymmetric dimethyl arginine (ADMA) levels is known to induce hypoxia driven mitochondrial dysfunction in the lungs, despite normoxic conditions. This alteration is known to be caused by interleukin-4 (IL-4). In addition to asthma, IL-4 and ADMA are also known to be elevated in metabolic syndrome (MetS), but the existence of hypoxic response in arterial tissue remains elusive. Since, MetS and asthma are correlated; we explored whether hypoxia exists in vascular tissue and is associated with IL-4 and ADMA levels.

To induce MetS, C57BL/6 mice were fed chow, high-fat or high-fructose diets for six months. Interestingly, MetS was associated with the induction of the hypoxic response in aortic tissue. Further, IL-4 and ADMA, which induce aberrant hypoxic response despite normoxia in the lungs, were elevated in the aorta of mice with MetS. Additionally, significant reductions in the levels of mitochondrial biogenesis factors (TFAM, PGC1α) and respiratory chain complexes (Complex I and Complex IV) activities were observed in MetS mice. Mitochondrial dysfunction in the aorta of mice with MetS perturbed mitochondrial inner membrane integrity and was associated with the leakage of cytochrome c in the cytosol. These results collectively suggest that elevated levels of IL-4 and ADMA in the aorta of mice with MetS correlates with induced vascular mitochondrial dysfunction and hypoxic response.

## Introduction

Atherosclerotic vascular disease and its clinical sequelae are the leading causes of morbidity and mortality in the Western world [1]. MetS has received considerable attention over the last decades as the cluster of its associated diseases significantly increase the risk for cardiovascular disease (CVD) mortality [2]; in fact, it is also known as the cardiometabolic syndrome. It is estimated that around a quarter of the world’s adult population have metabolic syndrome [3] and they are three times more likely to have a heart attack or stroke [4].

The mechanisms underlying cardiac dysfunction in the MetS are complex and might include lipid accumulation [5], increased fibrosis [6], altered calcium homeostasis [7], mitochondrial dysfunction [8] and increased oxidative stress (OS) [9]. Mitochondrial dysfunction contributes to cardiac dysfunction and myocyte injury by the loss of its metabolic capacity resulting in the release of toxic products [10]. Although, each component of the MetS causes cardiac dysfunction but in combination they carry additional risks [11,12]. Mitochondrial dysfunction is now considered as an important aspect of the pathogenesis of CVD. In addition, increased mitochondrial ROS can lead to cytochrome c release and the initiation of apoptotic cascade [13].

Of all the known contributing factors of CVD, one of the major important factors contributing to increased OS in CVD is hypoxia [14]. When tissue oxygenation decreases, the respiratory chain complexes become less active that lead to increased localized production of ROS in the mitochondrion, thereby, compromising ATP production and enhancing hypoxia [15,16]. This enhanced hypoxic condition, when coupled with mitochondrial dysfunction, may place aorta under bioenergetic and reductive stress, thereby contributing to the pathogenesis of CVD. In addition, in CVD, gene regulation promoted by a low oxygen microenvironment has received increased attention [17]. Further, the hypoxia-inducible miRNAs also termed as “hypoxamiRs” were found to play a cruicial role in CVD. miR-210 (HIF-1 transcriptional target) is a master hypoxamir [18], that controls cellular oxygen consumption and metabolism by regulating mitochondrial function [19,20]. Recent evidences have shown that intra-myocardial injection of non-viral vector expressing miR-210 precursor, stably expressed miR-210 for at least eight weeks in the heart [21], and improved cardiac function, reduced the infarct size and rectified angiogenesis after myocardial infarction [21], indicating miR-210 delivery as a therapeutic approach in ischemic heart disease.

In context of asthma, we have previously shown that IL-4, a marker of TH2 immune response, and ADMA, an analogue of L-arginine, together can induce hypoxic response and mitochondrial dysfunction in the normoxic lungs [22]. IL-4 promotes intracellular ADMA accumulation via altered expression of protein arginine methyltransferases (PRMT) and Dihydrodiamino hydrolases (DDAH) through calpain activation [22].

Therefore, hypoxic response [8] and ADMA [23] appears to be the important CVD risk factors, and their role in causing mitochondrial dysfunction in ‘asthma’ has been deciphered in one of the study from our group [22]. Moreover, large scale clinical trials using antioxidants therapies for the treatment of CVD have been disappointing because of the lack of efficacy and undesired side effects [24], we wanted to gain insights into the molecular events underlying the mitochondrial dysfunction in heart, an obligate aerobic organ. Here, we investigated the signalling mechanism underlying cardiac dysfunction in the MetS and found the role of IL-4 and ADMA driving pathological hypoxic response leading to CVD. We used high-fat (obese, HFA) and high-fructose (non-obese, HFR) diet induced mice models of MetS (C57BL/6 mice) [25] to show for the first time that diet-induced mitochondrial dysfunction in CVD is associated with hypoxic response, despite normoxic conditions.

## Material and Methods

### 1. Development of in-vivo model of MetS and diet plan

Four to five week old male C57BL/6 mice were obtained from National Institute of Nutrition (Hyderabad, India) and acclimatized for a week prior to starting the experiments. Animals were maintained as per the Committee for the Purpose of Control and Supervision of Experiments on Animals (CPCSEA) guidelines. All experimental protocols and methods were approved by the institutional animal ethics committee (IAEC; Protocol Number-03/IGIB/IAEC/MLP5502). Mice were divided into three groups (n = 6) and were named according to the diet provided as Control, HFA and HFR. The mice had free access to either a standard rodent chow (C) (5.5% fat and nil refined fructose), or a high-fat (HFA) diet having 60% of energy from fat, or a high fructose (HFR) diet with 70% energy from fructose. Mice were kept on this diet for eighteen weeks and sacrificed for the collection of aorta tissue.

### 2. IL-4 measurement

Whole aorta lysates serum samples from the MetS murine models were used for IL-4 ELISA. IL-4 measurement was done by using sandwich ELISA in which HRP-conjugated detection antibody was added. The substrate used for HRP was Tetramethylbenzidine (TMB) whose absorbance was measured at 450 nm as per the manufacturer’s protocol (BD Biosciences, San Deigo).

### 3. Measurement of ADMA levels

Whole aorta lysates serum samples from the MetS murine models were used for ADMA ELISA (DLD Diagnostika, Germany). Briefly, ADMA is bound to the solid phase of the microtiterplate; ADMA in the samples is acylated and competes with the solid phase bound ADMA for a fixed number of rabbit anti-ADMA antiserum binding sites. The antibody bound to the solid phase ADMA is detected by anti-rabbit/peroxidase. The substrate TMB/peroxidase was used. Results were expressed in nanomoles/microgram (nmoles/ug) protein.

### 4. Western blot analysis

Mitochondrion cytosol extraction protocol was followed using MEB (membrane extraction buffer) buffer (Thermo Fischer Scientific). Mitochondrion was collected in pellet and later saved in mitochondrion storage buffer however the cytosol was collected in supernatant. Whole aorta lysates and cytosols were separated by 10% SDS-PAGE, transferred onto PVDF membrane (MDI, India). Transferred membrane was blocked with blocking buffer (3%/5% Bovine Serum Albumin in PBS with Tween 20/ Skim milk/ NAP).The blots were then incubated with primary monoclonal antibody for HIF1α, TFAM, PGC1α and Cytochrome-c. Enhanced chemiluminescence Western blotting detection reagent (Pierce, Rockford, IL) was used to detect protein levels. Band densities of proteins were normalized to β-actin. Densitometry was performed by ImageJ software.

### 5. Immunohistochemistry

Formalin-fixed, paraffin-embedded aorta tissue sections were used for Immunohistochemistry (IHC). Commercial primary goat/rabbit polyclonal antibodies against HIF1α, TFAM, PGC1α and iNOS and their respective HRP conjugated secondary antibodies (Merck, USA) were used. IHC was performed as described previously [26].

### 6. cDNA synthesis and q-PCR

Total RNA was isolated from aorta tissue using TRIzol reagent (Invitrogen, Grand Island, NY, USA), quantified using a Nanodrop ND-1000 spectrophotometer. DNase (Ambion) treatment was given and 1µg of total RNA was used for cDNA synthesis. q-PCR was done for the expression analysis of miR-210-3p. U6snRNA used as an internal control for normalization. The detail of the primers is provided in **Supplementary Table 1.**

**Supplementary Table 1.**
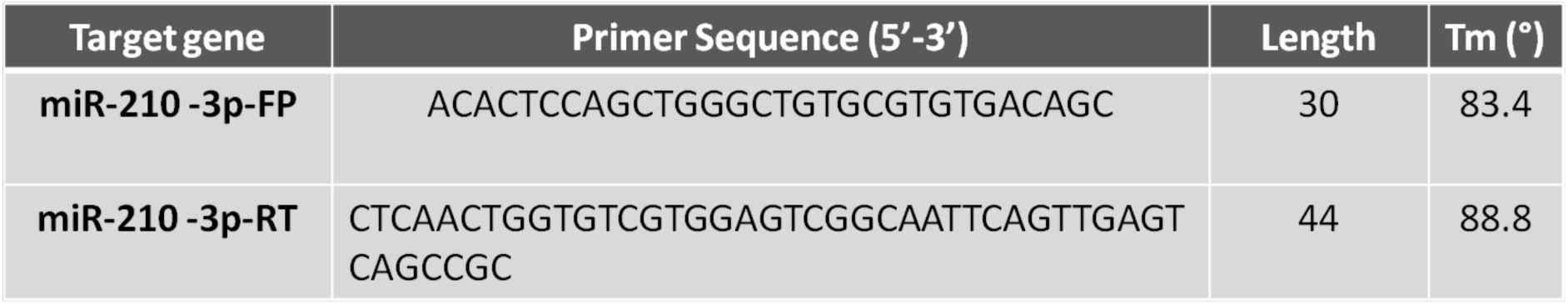
Primer sequences for miR-210-3p.

### 7. Measurement of Inner Mitochondrial Membrane

The mitochondria from whole aorta were isolated using mitochondrial isolation kit as per manufacturer’s instructions (SIGMA, USA). Inner Mitochondrial Membrane (IMM) of aorta mitochondria was determined by measuring the uptake of the cationic carbocyanine dye JC-1 (5,5’,6,6’-tetrachloro-1,1’,3,3’-tetraethylbenzimidazolcarbocyanine iodide) into the matrix. The fluorescence was read in spectrofluorometer and reported as fluorescence units in the mitochondria suspension per mg mitochondrial protein (FLU/mgP)

### 8. Measurement of mitochondrial Complex I and Complex IV activity

Complex I and Complex IV activities were measured in the isolated mitochondria from whole aorta fractions as per manufacturer’s protocol [27,28,29]. Briefly, COXETC (Complex IV) activity is based on the oxidation of ferrocytochrome c to ferricytochrome c by COXETC present in the mitochondria. Similarly, Complex I activity was determined following the oxidation of NADH to NAD+. 10 ug and 45 ug protein amount of aorta mitochondrial samples were used to measure Complex I and Complex IV activity, respectively.

### 9. Statistical analysis

Data are expressed as means ±SD. Differences between the groups were determined using Student’s t-test and P≤0.05 were considered significant. Graphs were prepared using Graph Pad Prism3 software.

## Results

### The characteristics of obese and non-obese model of MetS in mice

Aorta samples were taken from HFA and HFR diets fed mice, which were previously been used in a study [25] from our group for the characterization of lungs in MetS conditions **(Supplementary Table 2).** These mice had high levels of total mass & fat mass, blood pressure, cholesterol and triglycerides, increase in serum insulin as well as random blood glucose levels (published data) [25].

**Supplementary Table 2.** Metabolic features in the obese and non-obese model of MetS in mice. Total body mass, fat mass and blood pressure were significantly higher in both HFA and HFR groups in comparison to Control group with mice on normal diet. Both HFA and HFR diets were found to be associated with a significant increase in the blood glucose levels and high levels of serum cholesterol, triglycerides and insulin.

|  | Control | HFA | HFR |
| --- | --- | --- | --- |
| Total Mass (grams) | 28 | 52<sup>*</sup> | 32 |
| Fat Mass (grams) | 7 | 28 | 12 |
| Blood Pressure (mm Hg) | 140 | 150 | 220<sup>*</sup> |
| Cholesterol conc. (mg/ml) | 30 | 58<sup>*</sup> | 63<sup>*</sup> |
| Triglyceride Conc. (nmol/μl) | 0.4 | 0.75 | 0.74 |
| Glucose (mg/dl) | 201±58 | 383.33±26<sup>*</sup> | 451.33±46<sup>*</sup> |
| Insulin (ng/ml) | 0.32±0.04 | 0.67±0.10<sup>*</sup> | 0.44±0.03<sup>*</sup> |

### High-fat or high-sugar diets lead to increased IL-4 and ADMA levels with NO elevated levels

To determine whether the pulmonary features of elevated IL-4, ADMA and OS in MetS [25] **(Supplementary Table 2)** are also present in vascular tissue, we measured these parameters in the aorta of Control (C) mice and in the mice fed with HFA and HFR of MetS groups. IL-4 levels were found to be significantly elevated by ∼1.87 folds and ∼2.13 folds in the aorta of HFA and HFR groups, respectively **(Fig 1a).** Similarly, ADMA levels were found to be significantly increased by ∼2.26 folds and ∼3.34 folds in the aorta of mice fed with HFA and HFR diet, respectively, compared to C group **(Fig 1b).** We next investigated, whether the levels of inducible nitric oxide synthase (iNOS) enzyme that regulates NO production were differentially expressed in aorta of differently fed MetS mice. Western blot analysis showed that iNOS expression were trending to slight higher levels in both HFA and HFR MetS mice groups (∼1.4 and 1.23 folds, respectively) in comparison to C group **(Fig 1c)**. Importantly, similar trends were seen in the IHC analysis of iNOS **(Fig 1d).**

**Fig 1:**
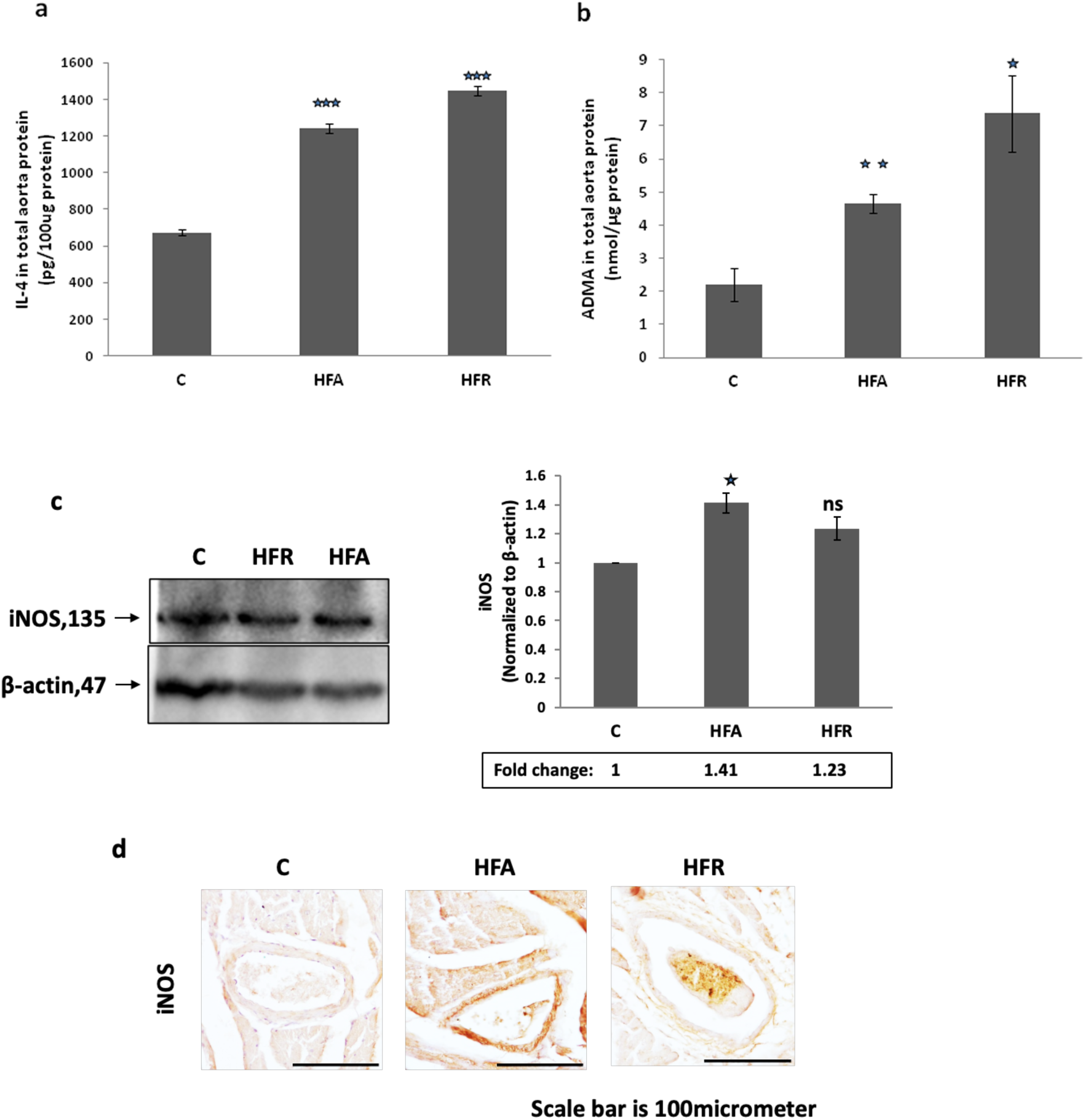
Induction of IL-4, ADMA and iNOS in the aorta of mice with MetS. **(a,b)** IL-4 and ADMA levels were estimated from C, HFA and HFR mice (n=6) in whole aorta lysates using ELISA. Increased IL-4 and ADMA levels were measured in total aorta lysates of HFA and HFR mice. Data represents mean ± SD; n=6; *p ≤ 0.05, **p ≤ 0.01 or ***p ≤ 0.001 was considered to be statistically significant; ns-non significant. **(c)** Western blot analysis of iNOS **(d)** Representative IHC images (n=3) of iNOS in the aorta of mice. Brown colour indicates the positive expressions. Data represents mean ± SD;*p ≤ 0.05, **p ≤ 0.01 or ***p ≤ 0.001. Images are at 20X objective magnification. Here, C is for Control, HFA is for High Fat diet, HFR is for High Fructose diet.

**Supplementary Table 3.** Levels of IL-4, ADMA and Inducible Nitric oxide synthase (iNOS) in the lungs of mice with MetS. Increased levels of IL-4 and ADMA with induced expression of iNOS were observed in whole lung lysate of HFA/HFR mice as compared to Control mice. Data shown here are Mean ± SE of 6 mice in each group.*Denotes statistically significant differences (p < 0.05) vs. Control.

|  | Control | HFA | HFR |
| --- | --- | --- | --- |
| IL-4 protein Conc. in whole lung (pg/100ug protein) | 17 | 27<sup>★</sup> | 23<sup>★</sup> |
| ADMA protein Conc. in whole lung (nmol/ug protein) | 0.9 | 2<sup>★</sup> | 3.8<sup>★</sup> |
| iNOS protein levels in whole lung (fold change) | 1 | 2.2<sup>★</sup> | 1.7<sup>★</sup> |

### Intracellular ADMA accumulation in MetS upregulates HIF1α and induces miR-210 expression in the aorta

Further, to determine whether the pulmonary features of elevated HIF1α are also present in vascular tissue, we measured HIF1α expression levels in the aorta of mice with diet induced MetS (HFR and HFA diet fed mice). In Western blot **(Fig 2a)** and ELISA **(Fig 2b)** analysis, HIF1α expression levels were found to be significantly up-regulated (∼ 1.5-2.5 folds) in the aorta of HFA and HFR mice, respectively. Importantly, similar trends were seen in the IHC analysis of HIF1α in aorta **(Fig 2c).**

**Fig 2:**
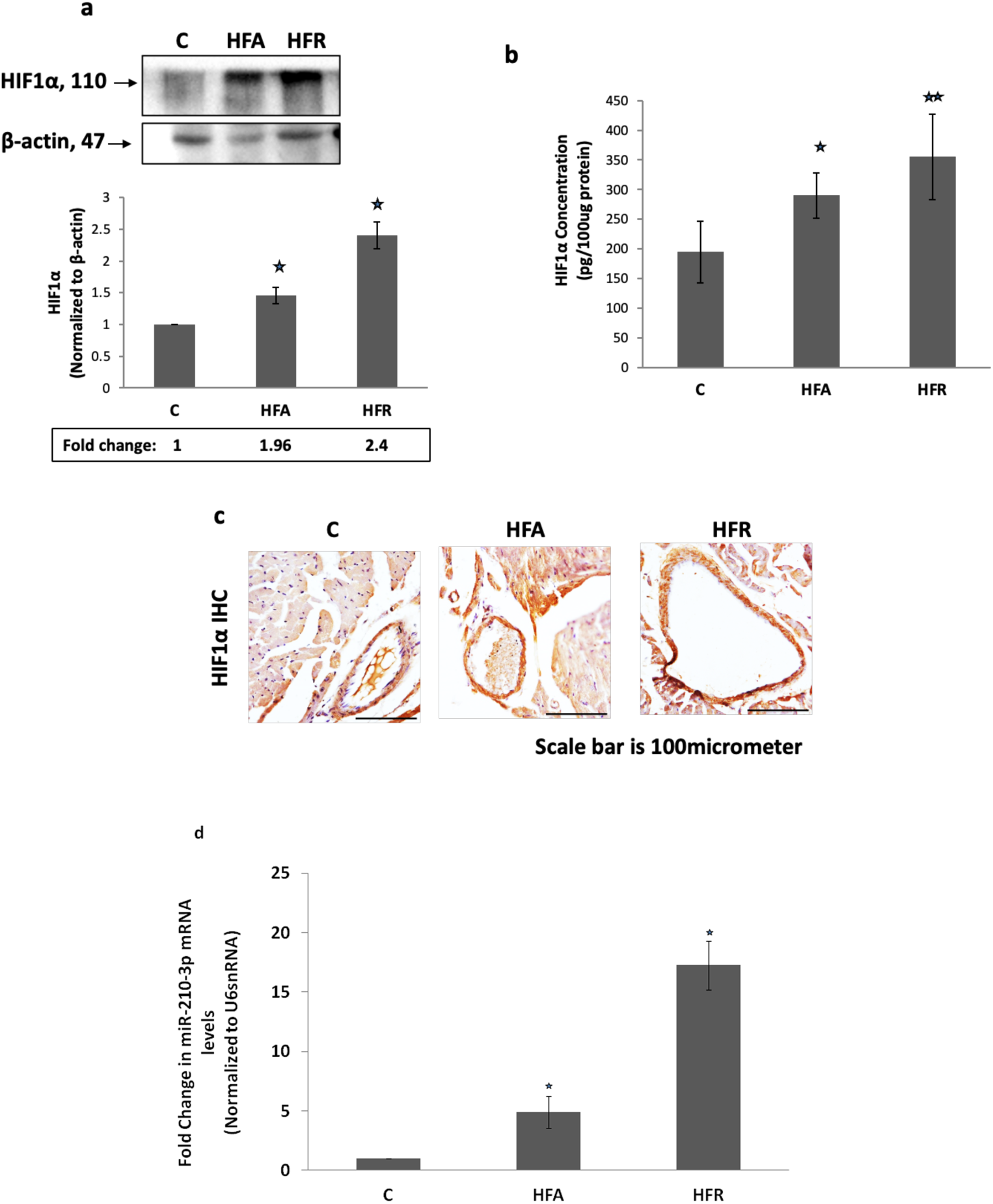
Increased HIF1α level and miR-210 expression in aorta of mice with MetS. **(a, b)** Western blot analysis (a) and ELISA (b) of HIF1α in HFA and HFR MetS mice groups **(c)**Representative IHC images of HIF1α in the aorta of mice. Brown colour in IHC images indicates the positive expressions. Images are at 20X objective magnification. Data represents mean ± SD; n=6; *p<0.05, **p ≤ 0.01. **(d)** Real Time PCR of miR-210. Experiment was done in triplicates. Each value is mean ± SD; n=3; *p<0.05. Here, C is for Control, HFA is for High Fat diet, HFR is for High Fructose diet.

In addition, to confirm hypoxia signalling in the aorta of MetS mice model, we measured the expression of microRNA-210 (miR-210), a direct downstream target of HIF1α. miR-210 expression was found to be multi-fold up-regulated, by 5 folds and 17.3 folds in HFA and HFR mice groups, respectively, in comparison to C group **(Fig 2d).**

### Increased hypoxic response correlates with the decreased expression of mitochondrial turnover proteins (PGC1α, TFAM) in the aorta of mice with MetS

To determine whether the elevated cellular hypoxic response observed in aorta of MetS mice led to any changes in the mitochondrial turnover, we measured the expression of PGC1α and TFAM proteins in aorta of mice with MetS. Western blot analysis of the HFA or HFR induced MetS mice group showed reduced expression of PGC1α, (-by ∼30%) **(Fig 3a)** and TFAM (-by∼32%) **(Fig 3c)**.Similar trends were observed in IHC analysis of PGC1α and TFAM in aorta tissue **(Fig 3b and 3d).**

**Fig 3:**
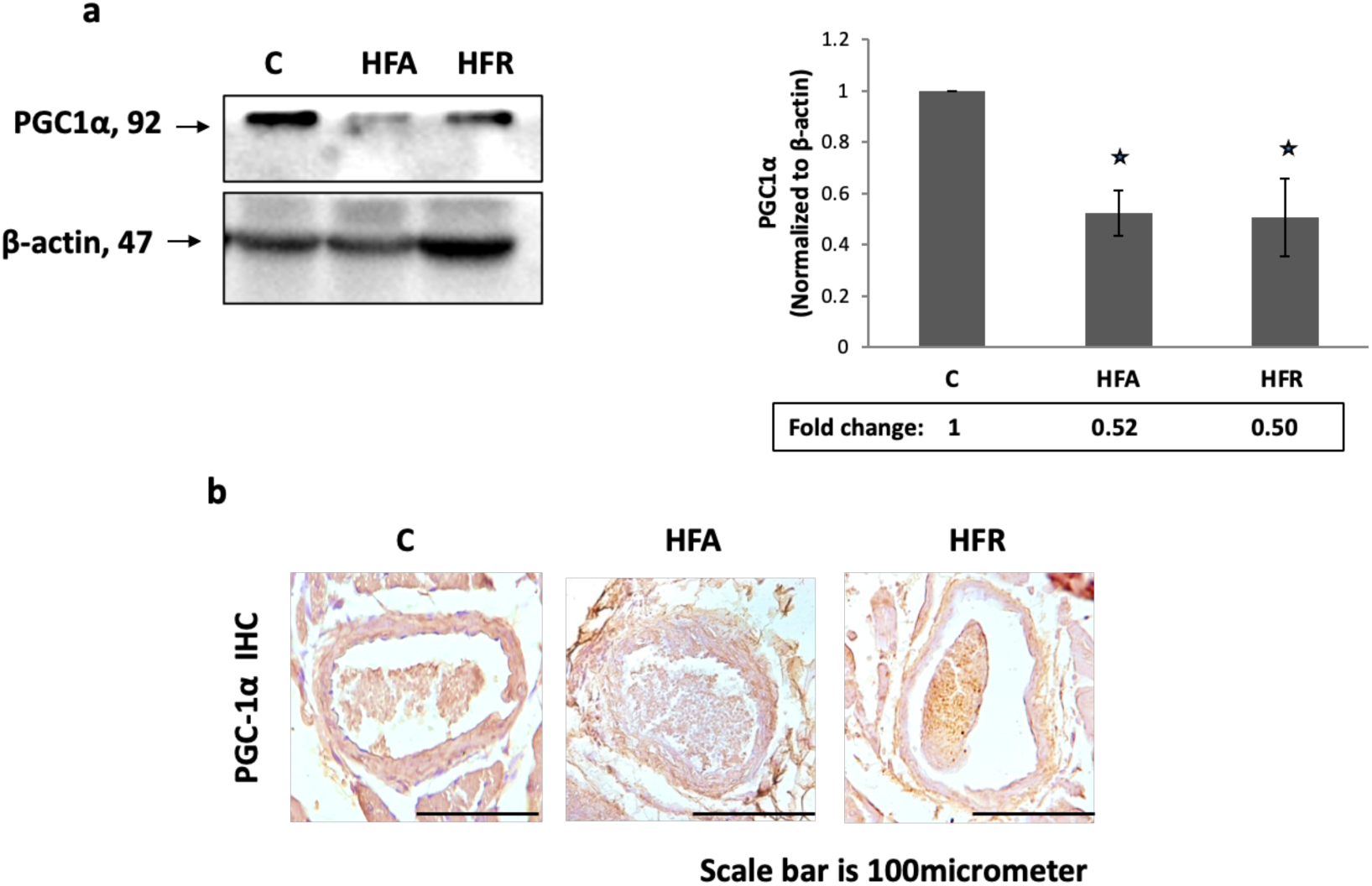
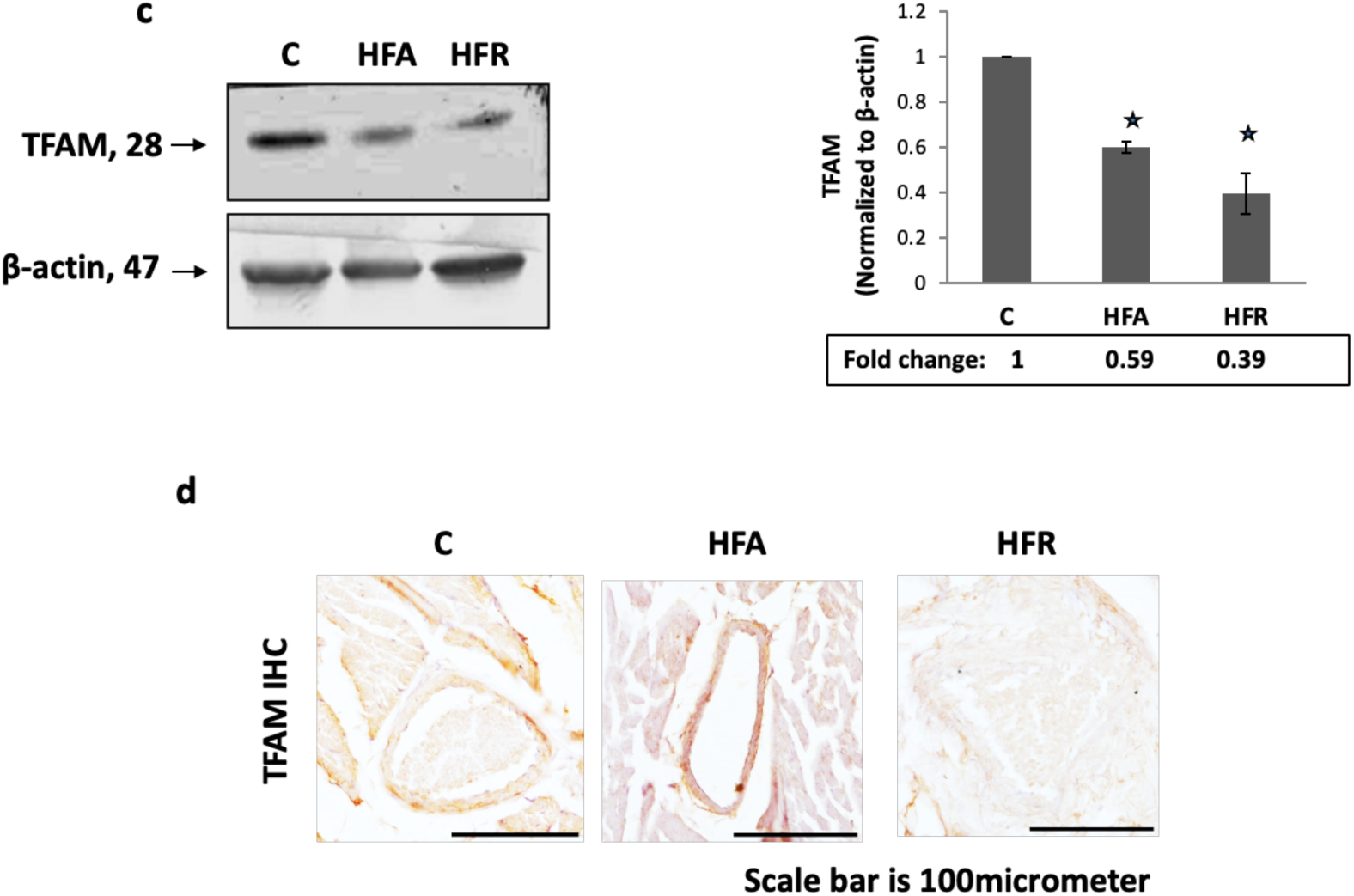
Decrease in PGC1α and TFAM levels in the aorta of mice with MetS. **(a)**Western blot analysis of PGC1α in whole aorta lysates of HFA and HFR MetS mice groups; n=6 **(b)** Representative IHC images of PGC1α. Brown colour indicates positive expression. Images are at 20X magnification. **(c)** Western blot analysis of TFAM in HFA and HFR MetS mice groups; n=6 Each value is mean ± S.D. *Denotes statistically significant differences (P<0.05) vs. Control; ns-non significant. **(d)** Representative IHC images of TFAM. Brown colour indicates the positive expressions. Images are at 20X objective magnification. Here, C is for Control, HFA is for High Fat diet, HFR is for High Fructose diet.

### Increased hypoxic response led to reduced mitochondrial function in mice with MetS

Lastly, to determine whether slight elevated NO levels driven hypoxic response in mice with MetS correlates with mitochondrial dysfunction, thereby contributes to CVD, we first measured mitochondrial Complex I and IV activities in the aorta of MetS mice. Interestingly, HFA and HFR MetS mice groups showed significant reductions in the mitochondrial complex I **(Fig 4a)** and complex IV activities **(Fig 4b).** To examine, whether decreased activity of Cytochrome c oxidase (COX_ETC_) disrupts mitochondrial inner membrane integrity (IMM), we measured mitochondrial membrane potential (ΔΨm) using JC1 dye in the isolated mitochondria from the aorta. As expected, we found significant reductions in ΔΨm in the mitochondria from the aorta of both HFA and HFR MetS mice **(Fig 4c)**. As COX_ETC_ activity was found to be reduced in the aorta of MetS mice, and NO production causes the mitochondrial pore formation, we analyzed the cytochrome c levels in the aorta cytosol. The levels of cytochrome c were significantly increased (1.5-3 folds) in the cytosols of HFA and HFR fed cardiometabolic mice aorta, compared to normal control mice **(Fig 5)**. Thus, these results collectively evidence the proof of mitochondrial dysfunction in the context of MetS murine models.

**Fig 4:**
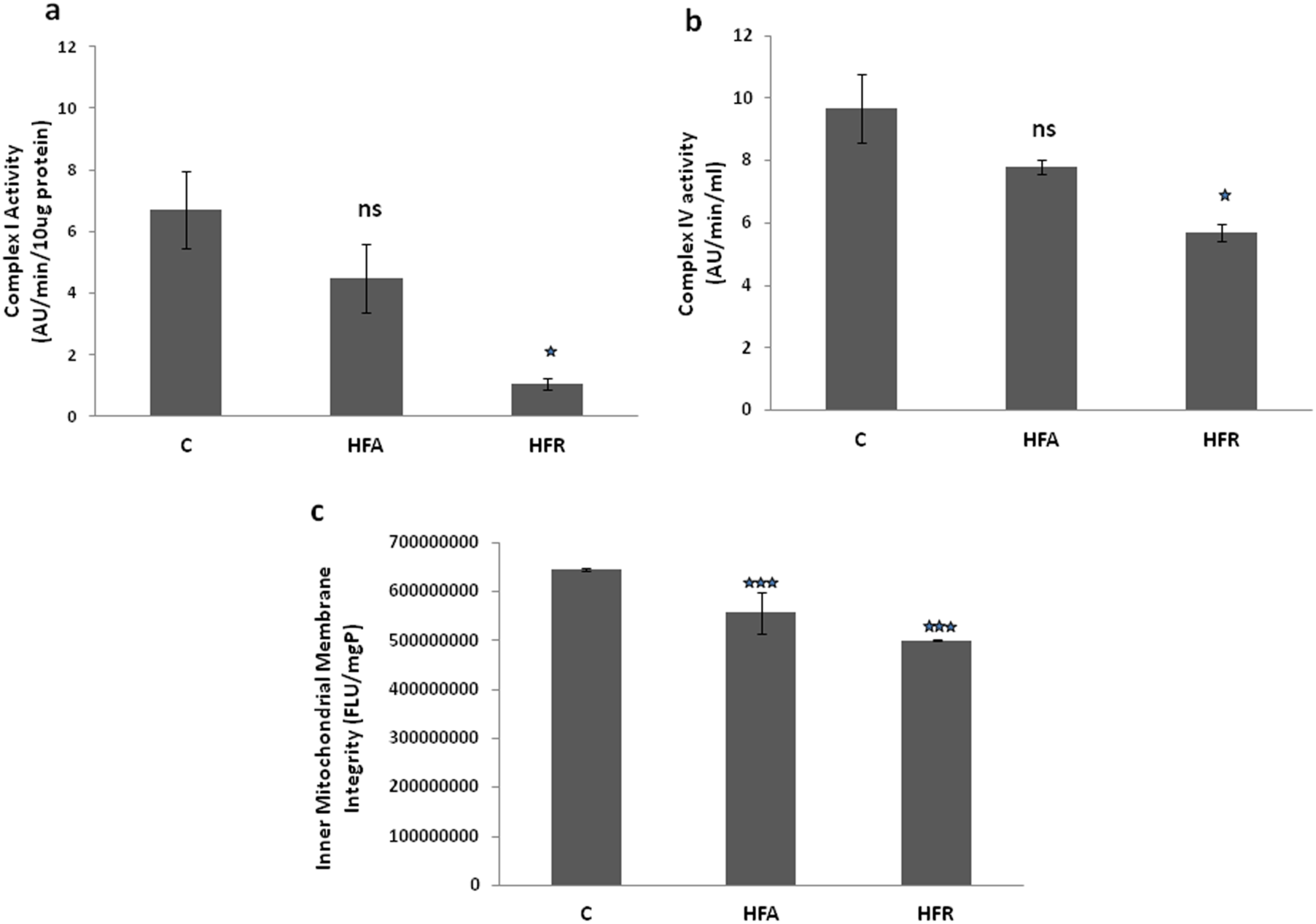
Altered mitochondrial functions in aorta of mice with MetS. **(a,b,c)** Mitochondrial Complex I and complex IV activities and Integrity of inner mitochondrial membrane were determined in the isolated aorta mitochondria of MetS. Data represents mean ± S.D.; n=8; *p ≤ 0.05, **p ≤ 0.01 or ***p ≤ 0.001was considered to be statistically significant; ns-non significant. Here, C is for Control, HFA is for High Fat diet, HFR is for High Fructose diet.

**Fig 5:**
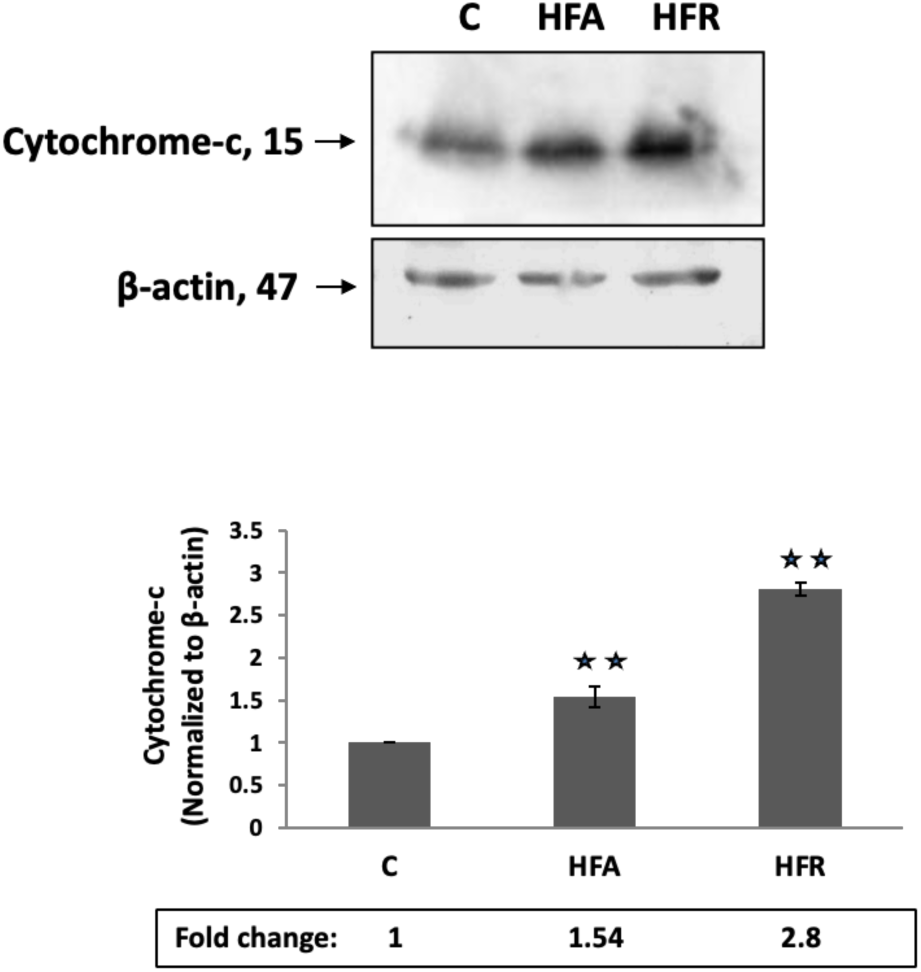
Leakage of mitochondrial cytochrome-c to cytosol in the aorta of mice with MetS. Cytochrome c levels were measured in cytosolic fractions of aorta. Data represents mean ± SD; n=4; *p <0.05, **p ≤ 0.01 or ***p ≤ 0.001 was considered to be statistically significant. Here, C is for Control, HFA is for High Fat diet, HFR is for High Fructose diet. Cytochrome-c blot was developed using ECL (enhanced *chemiluminescence*); however, β-actin was developed using DAB (3, 3’-diaminobenzidine) substrate.

## Discussion

In the present study, we addressed the hypothesis that whether chronic exposure to nutritional insults, either HFA or HFR, leads to metabolic induction of a hypoxic response in vascular tissue, despite normoxic conditions. For this, we manipulated the diets of C57BL/6 mice to induce MetS, independent of obesity or adiposity. A high fat diet in these mice was associated with obesity, adiposity, hyperlipidaemia and hyperglycaemia; while a high fructose diet led to hyperlipidaemia, hyperglycaemia, and hypertension, but no obesity or adiposity **(Supplementary Table 1)**. As previously published, HFA and HFR diets were found to be associated with MetS features [25].These extreme diets mimic “junk foods” and induce a MetS. Our main hypothesis here was centred on the potential role of the transcription factor HIF1α, as a link between MetS and CVD, via mitochondrial dysfunction.

This study, for the first time, demonstrated that the observed mitochondrial dysfunction in cardiometabolic syndrome was associated with the expression of HIF1α in the aorta, which likely could be driven by abnormal Arginine-NO metabolism, as suggested by Williams D et al, 2015, Morris SM Jr, 2005 [26,30] and indicated by our data; NO and ADMA levels.

A number of previous studies have demonstrated that the pro-oxidative and pro-inflammatory pathways within vascular endothelium play an important role in the initiation and progression of atherosclerosis [31–33]. In addition, a recent study has provided compelling evidence to indicate that IL-4, a marker of Th2 immune response, can induce pro-inflammatory environment via oxidative stress-mediated up-regulation of inflammatory mediators such as cytokine and chemokine in vascular endothelial cells [34]. IL-4 is often seen to be upregulated in subjects with MetS [35–36] and we have previously found IL-4 to be potentiating ADMA accumulation in airway epithelial cells, via induction of protein arginine methyltransferases (PRMTs) [22]. Here, we observed an increase in IL-4 levels in aorta of mice with HFA and HFR diets. ADMA, an analogue of L-arginine, is a naturally occurring product of arginine metabolism found in human circulation (0.45 ± 0.19 μmol/L). ADMA has been demonstrated to be not only a marker of endothelial dysfunction [37], but also a novel cardiovascular risk factor [23]. Here, we found significant increase in levels of ADMA in aorta of mice fed with HFA or HFR diets. This is the first report tying together IL-4 and ADMA increase in vascular tissue of mice with diet-induced metabolic syndrome.

It is known that elevated levels of ADMA inhibit NO synthesis in endothelial cells and therefore impair endothelial function and thus promote atherosclerosis [38–40]. Moreover, experimental and clinical evidences have shown that ADMA significantly contributes to oxo-nitrative stress via the generation of superoxide anions (O2−) by uncoupled nitric oxide synthase, which reacts with NO in the endothelial cells leading to the formation of peroxynitrite, a highly reactive oxidant species [41]. We thus hypothesized that increased levels of ADMA may be responsible for oxo-nitrative stress and organelle dysfunction leading to CVD. So, we further investigated whether enzyme that regulates NO production i.e. iNOS was differentially expressed in aorta of differently fed MetS mice. The levels of iNOS which we are showing here, as a major determinant of ADMA induced hypoxia, are not significantly consistent among HFA and HFR groups. Possibly, there could be mechanisms independent of iNOS for ADMA effects on hypoxia, in concordance with previously reported studies [42]. This could be an explanation for not finding the major changes in the levels of iNOS. Therefore, in our study the pathophysiological concentrations of ADMA can regulate hypoxic gene expression in MetS murine endothelial cells by what appears to be an NO-independent mechanism.

To further elucidate the mechanism of IL-4 driven ADMA mediated increase in hypoxic response, despite normoxic conditions in aorta, similar to what would be expected in well oxygenated arterial tissues, we measured HIF1α expression in the aorta of mice with diet induced metabolic syndrome and found significant increase in HIF1α in both HFA and HFR groups. We further determined the expression levels of microRNA-210 (miR-210), a direct downstream target of HIF1α, to confirm that the cellular hypoxic response was functionally active. As expected, miR-210 expression was found to be markedly high in the aorta of mice with MetS, indicating a strong possibility of functionally active cellular hypoxic response in vascular tissue of such mice. miR-210 has been implicated in several key aspects of cardiovascular diseases and cancer [18,43,44], and miR-210 overexpression has been seen to be responsible for mitochondria dysfunction [45]. We speculate that this may be a key mechanism linking the hypoxic response to systemic biochemical changes of MetS as described here to local mitochondrial dysfunction in endothelial cells.

Mitochondria are known as the “powerhouse” of the cell, generating ATP via oxidative phosphorylation (OXPHOS) complexes, including Complexes I-IV, which are present in the inner membrane of mitochondria [46]. To determine whether hypoxic response was associated with mitochondrial dysfunction in our model, we checked activities of mitochondrial complex I and IV in the aorta of MetS mice groups, where we had observed enhanced hypoxic response. The results showed marked reductions in both mitochondrial complex I (NADH dehydrogenase) and IV (cytochrome c oxidase) activities in aorta of HFA and HFR MetS mice. Further, Cytochrome c oxidase of electron transport chain COX_ETC_ is an integral enzyme of inner mitochondrial membrane and decreased activity of COXETC causes reduction of IMM integrity [47]. To test this, IMM integrity for all groups was measured by the uptake of membrane potential sensitive dye: JC-1, which was found to be significantly decreased in the aorta of both HFA and HFR mice groups. There are many studies which suggest that oxidative stress (ROS and RNS) in mitochondria combines with peroxynitrite formation and triggers mitochondrial pore formation, thereby leading to cytochrome c leakage [48–49]. Since peroxynitrite also causes inhibition of cytochrome c oxidase and in our study COXETC activity was found to be reduced in the aorta of MetS mice, we analyzed the levels of cytochrome c in cytosol of aorta of MetS. The results demonstrated increased leakage of cytochrome c in the cytosols from aorta of HFA, HFR MetS groups. We also investigated whether mitochondrial biogenesis may be impaired. PGC1α acts as a co-activator in driving the expression of TFAM gene, which further translocates to nucleus and mediates mitochondrial transcription initiation and replication [50–52]. Both PGC1α and TFAM levels were found to be reduced in the aorta of HFA and HFR mice. Thus, the increased hypoxic response in MetS mice aorta was found to be associated with decreased expression of mitochondrial turnover proteins. These results together suggest the role of hypoxic response in causing mitochondrial dysfunction and decreased turnover, which may lead to CVD.

Putting these disparate evidences together, we speculate endothelial aorta is susceptible to ADMA induced hypoxic response and that this drives cardiovascular disease in metabolic models of CVD via mitochondrial effects. There is obviously no hypoxemia in the aorta of such mice and this is entirely a metabolically triggered cellular hypoxic response in normoxic conditions, as described by us previously in context of asthma [22]. Exposure to IL-4 further sensitizes endothelial aortic tissue to ADMA exposure. Taken together, our results suggest that intracellular ADMA accumulation, along with IL-4 causes mitochondrial dysfunction, cellular injury and tissue remodelling (Fig 6). We speculate that this contributes to the pathogenesis of CVD in diet induced metabolic syndrome.

**Fig 6:**
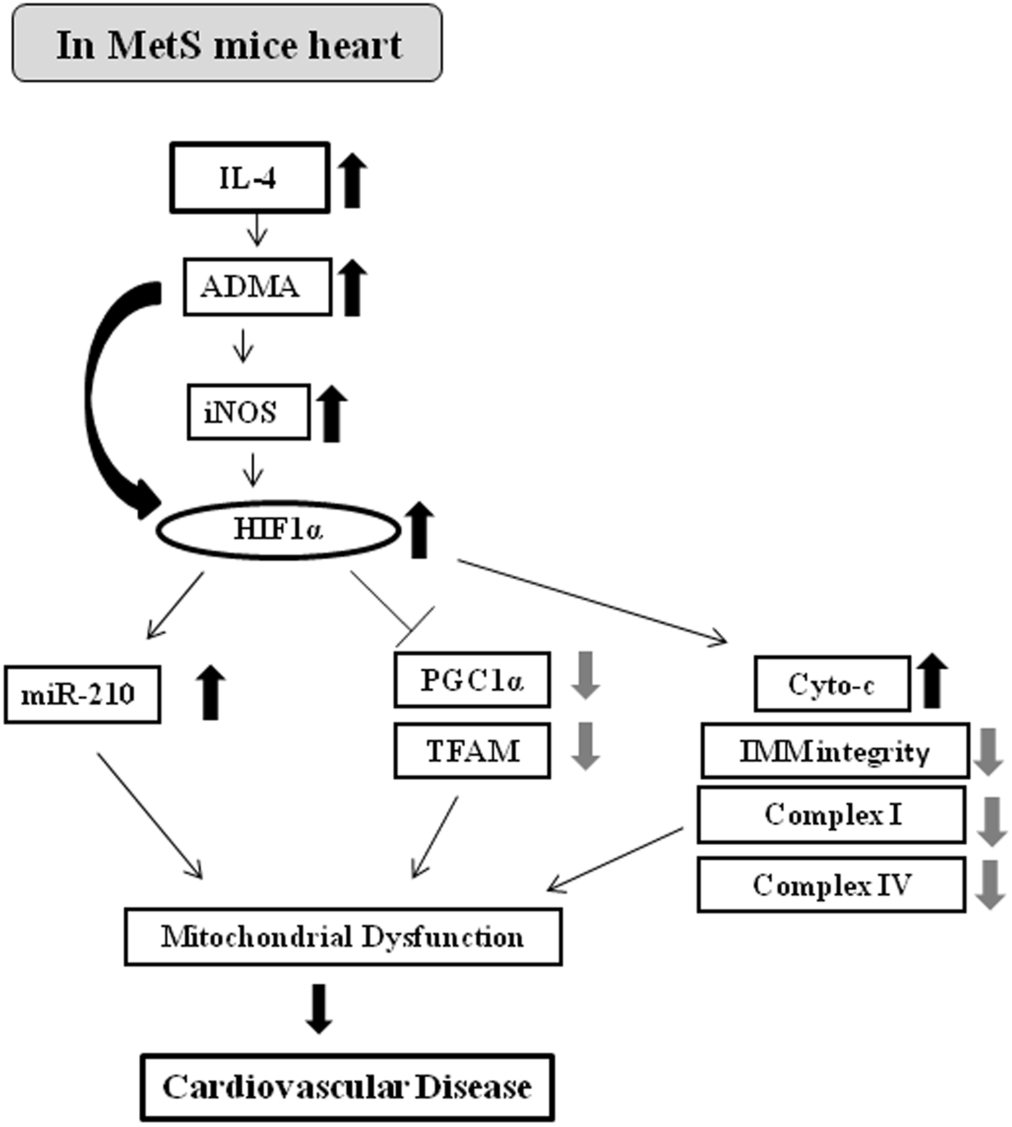
Proposed schematic diagram of ADMA-HIF1α axis affecting mitochondrial function and leading to CVD in MetS mice aorta.

## Conclusion

In this study, we wanted to gain insights into the molecular events underlying the mitochondrial dysfunction in aorta, an obligate aerobic organ. Here, we investigated the signalling mechanism underlying cardiac dysfunction in the MetS and found the role of IL-4 and ADMA driving pathological hypoxic response in leading to CVD. For this, we have used high-fat (obese, HFA) and high-fructose (non-obese, HFR) diet induced mice models of MetS (C57BL/6 mice). We demonstrated that the observed mitochondrial dysfunction in cardiometabolic syndrome was associated with the expression of HIF1α in the aorta, which likely could be driven by abnormal Arginine-NO metabolism. We show for the first time that diet-induced mitochondrial dysfunction in cardiovascular disease is associated with hypoxic response, despite normoxic conditions.

## Acknowledgements

This work was funded by grant from Council of Scientific and Industrial Research (CSIR, India) to AA. Research fellowship to EM is from Council of Scientific and Industrial Research (CSIR, India). VJ received research fellowship from DBT, Delhi, India. We would like to thank Dr. Anurag Agarwal and Dr. Vijay Pal Singh, IGIB, for providing MetS mice aorta samples and all the assistance pertaining to this work. The help by Dr. U. Mabalirajan, Bijay Ranjan Pattnaik, Naveen Kumar Bhatraju and other lab members is highly acknowledged. This work was supported by project MLP5502 (CSIR).

## Authors Contributions

EM designed experiments. EM performed experiments. VJ assisted with analysing IHC slides and data. VPS provided MetS mice aorta samples. EM wrote the manuscript. EM and VJ revised the manuscript.

## Footnotes

1 Evanka Madan: Sir Ganga Ram Hospital

